# Integrating experimental data to calibrate quantitative cancer models

**DOI:** 10.1101/032102

**Authors:** Heiko Enderling

## Abstract

For quantitative cancer models to be meaningful and interpretable the number of unknown parameters must be kept minimal. Experimental data can be utilized to calibrate model dynamics rates or rate constants. Proper integration of experimental data, however, depends on the chosen theoretical framework. Using live imaging of cell proliferation as an example, we show how to derive cell cycle distributions in agent-based models and averaged proliferation rates in differential equation models. We focus on a tumor hierarchy of cancer stem and progenitor non-stem cancer cells.

## I. Proliferation rate in cancer models

Simulation of tumor growth includes the crucial process of cell proliferation. Simulations with faster proliferation rates yield faster growing tumors. It is imperative that proliferation is properly calibrated, especially for quantitative models in which timing is crucial, such as fixation of mutations in a population, fractionated radiotherapy, or survival rates.

## II. Experimental proliferation rate

A cell’s potential doubling time may be derived from *in vitro* experiments, including live cell imaging. The fate of individual cells is followed, and the time interval between successive mitosis events scored. For the U87-MG human glioblastoma cell line, such analysis of 19 mitotic events yielded an average doubling time of 26±4 hours [1] (Fig. 1).

**Figure 1.**
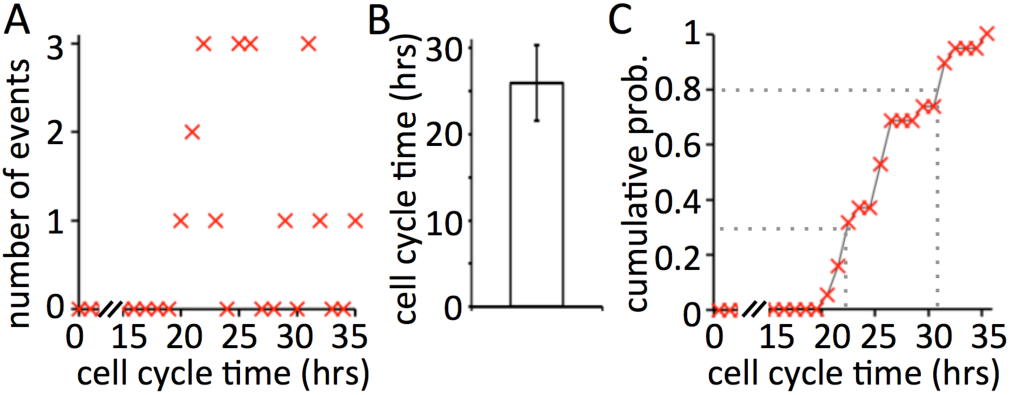
Experimentally measured doubling time of the U87-MG cell line (n=19). A: discrete recording of occurences of different cell cycle times. B: Average cell cycle time of of 26±4 hours. C: Cumultative density function for the distribution of cell cycle times. Dotted lines show probabilities of two cell cycle times (30% → 22.5hrs; 80% → 30.5hrs)

## III. Calibrating agent-based models

In stochastic agent-based models with individual cell resolution, each cell acts independently and thus needs to be assigned their individual doubling time. On the population level, the average cell cycle time should mimic T_pot_, but biological deviation must be reproduced. The probability of individual proliferation events must be scaled according to the simulation time, which must be less or equal to the unit of the most frequent events. In macroscopic or population-level simulations, this may be the unit of T_pot_ (hours or days); in microscopic simulations this might be cell migration (minutes) or diffusion of chemicals (seconds).

Let us consider non-spatial tumor growth with a simulation time step of Δ*t*=1 hour. Then, each *in silico* U87-MG cell should divide on average every 26 simulation time steps. Applying a uniform distribution to the average cell cycle time observed experimentally (i.e., each *in silico* cell has a 1/26 probability to divide in each time step) gives the correct average proliferation rate but a more than four times larger standard deviation. With a uniform distribution, 3.8% of *in silico* cells may divide already at the first time step. Similarly, some cells may stochastically not divide for more than 100 time steps (Fig. 2A). A uniform distribution applied to the cumulative probability derived from the experimental data (c.f. Fig. 1C) yields the correct average proliferation rate and standard deviation (Fig. 2B). Many agent-based models are calibrated from average data reported in the literature. Caution is warranted to use averages without high-resolution information about the actual distribution of events, especially when simulating rare stochastic events [2].

**Figure 2.**
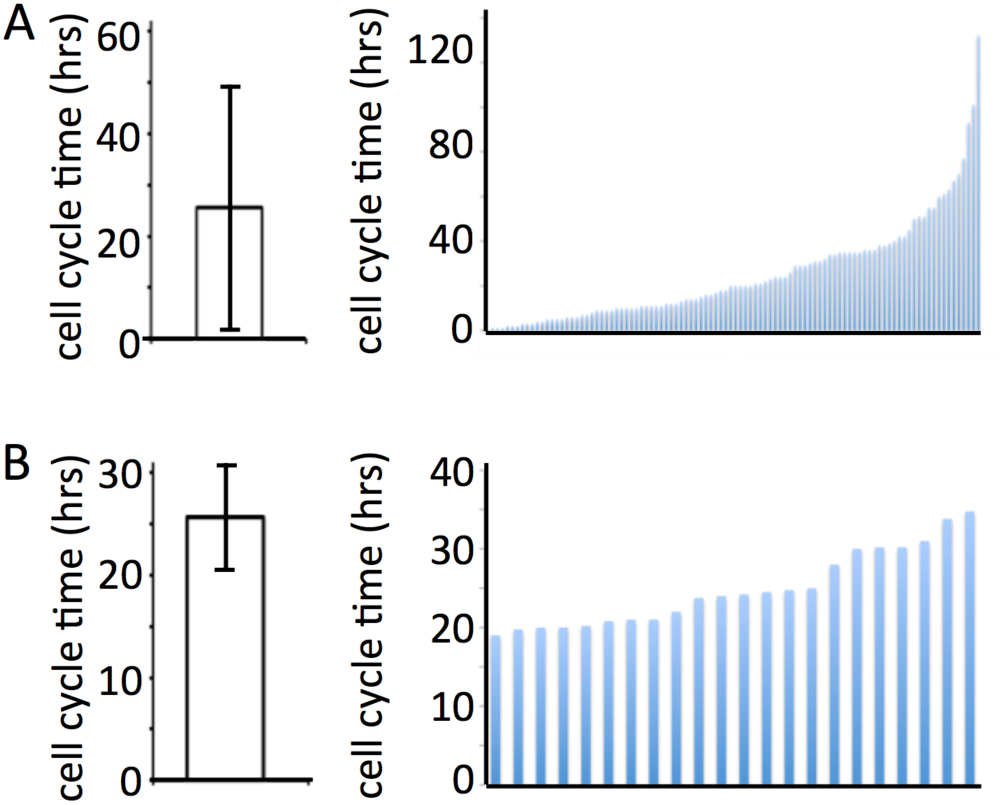
Simulated cell doubling times of *in silico* U87-MG cells. A: Uniform distribution applied to average cell cycle doubling time (left panel), and individual cell cycle time for 100 independent *in silico* cells. B: Uniform distribution applied to cumulative probability (left panel; c.f. Fig. 1B,C), and individual cell cycle times for 22 independent *in silico* cells.

## IV. Calibrating differential equation models

A differential equation of tumor growth includes a growth term – often either exponential or sigmoidal growth [3]–[5] – with a growth rate per unit time parameter, for demonstration purposes arbitrarily denoted λ. For exponential growth, the growth rate λ=ln 2/*T_pot_*, where *T_pot_* is the potential doubling time [6].

For illustration purpose we discuss the multicompartment ordinary differential equation (ODE) model of cancer stem cell-driven solid tumor growth by Weekes et al. [7], which is build after an agent-based model [8]. In brief, the population of cancer stem cells, *C(t)*, gives rise to *m* generations of non-stem progenitor cells *N_i_*, *1*≤*i*≤*m*. A cell in the *i*^th^ compartment leaves this compartment upon proliferation, and two new cells enter the *i*+*1*^st^ compartment. Cells in the *m*^th^ compartment die upon proliferation attempt.

Cancer stem cells divide symmetrically with probability *p_1_*, asymmetrically with probability *p_2_*, and may commit both daughter cells into differentiation with probability *p_3_ = 1−p_1_ − p_2_*. Both stem and non-stem cancer cells may die spontaneously, denoted by parameters *a* and *b*, respectively. The proliferation rate λ is identical and constant for cells in all compartments, which gives the equation system (other parameters and model details in [7])

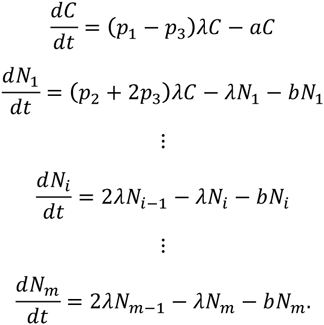

The dynamics of this multicompartment model mimics the population dynamics of a non-spatial agent-based model of cancer stem cell driven solid tumor growth. Under the specific assumptions detailed in [7], all non-stem cancer cells can be combined into a single compartment *H*

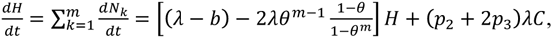

which, together with the equation for the cancer stem cell population, we call the derived two-compartment model. The derived two-compartment model is in good agreement with the dynamics of the multicompartment model (Fig. 3A).

**Figure 3.**
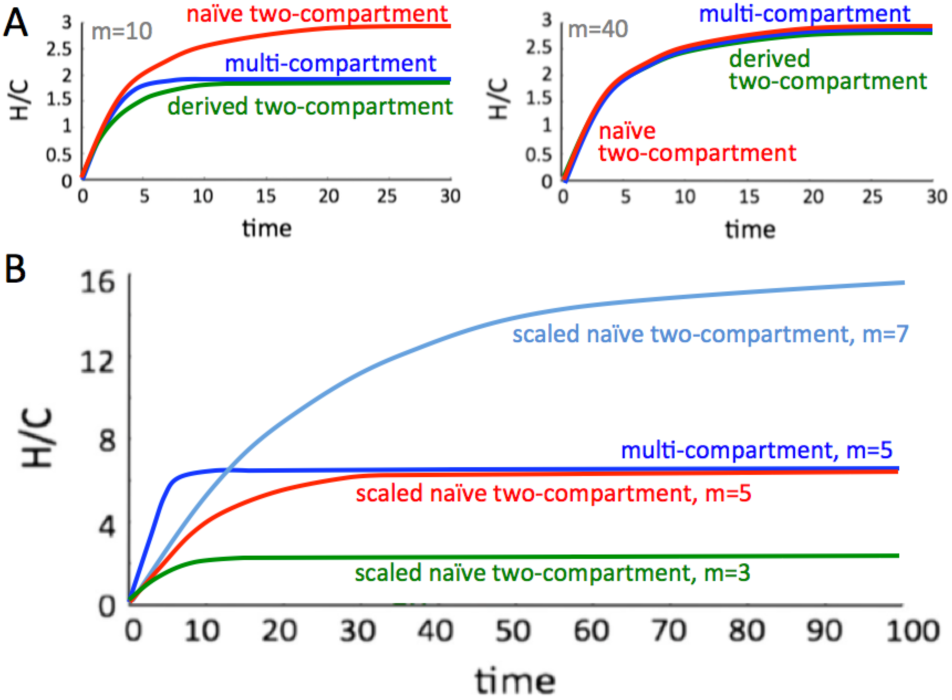
Time-dependent number of non-stem cancer cells (H) per cancer stem cell (C). A: The naïve two-compartment model assuming infinite number of generations, *m*→∞ performs poorly for low (left panel, *m*=10) but increasingly better for large *m* (right panel, *m*=40). The derived two-compartment model approaches the multi-compartment long-term outcome. B: Generation-adjusted non-stem cell compartment growth rate for different generations (*m*=3,5,7), copared to a multi-compartmental model with 5 compartments. Details and paramters in [7].

Assuming an infinite number of compartments m→∞ and other specific conditions detailed in [7], this model reduces to the linear two-compartment ODE model as developed by Hillen et al. [9] and others with the simplified general form

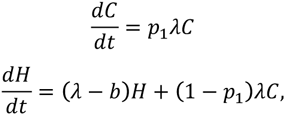

which because of its intuitive deviation we call the naïve two-compartment model.

As in the agent-based model, the cells in all compartments are assigned the same growth rate λ. As expected from model development assumptions, the naïve two-compartment model performs satisfactory for large generational hierarchy depths *m*, but poorly for smaller *m* (Fig. 3A). To account for the transitions between compartments and cells leaving the terminal compartment, the proliferation rate in the naïve two-compartment model needs to be scaled with 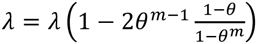, with 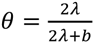 as discussed in detail in [7]. The scaled naïve two-compartment model with explicit consideration of the generational hierarchy depth *m* and terminal cell death *b* in the non-stem population growth rate enables adequate approximation of the multicompartmental model long-term behavior (Fig. 3B).

## V. Discussion

It becomes increasingly important for quantitative cancer modeling to calibrate model parameters; however, the integration of experimental data into model parameterization is not straightforward. Models must not only reproduce average behavior but also biological standard deviation to become meaningful. To this extend, the actual distribution of biological rates must be quantified and reported, which is rarely an observable in qualitative experimental setups. Whilst mathematical modeling could help simulate tumor growth on basic assumptions, the outcomes often depend heavily on chosen model parameters. Herein we have shown that the proliferation rate, which is the first term in most quantitative models, may not be as intuitive to integrate into a theoretical model. Upmost care must be taken when designing models as well as calibrating and validating them with experimental data. Understanding the limitations in data and modeling results is paramount in proper utilization of quantitative models in cancer research.

## Acknowledgment

H.E. would like to thank Jan Poleszczuk for critical reading of this article.

